# Temporal Trends without Seasonal Effects on Gestational Diabetes Incidence Relate to Reductions in Indices of Insulin Secretion: The Cambridge Baby Growth Study

**DOI:** 10.1101/556886

**Authors:** Clive J. Petry, Benjamin G. Fisher, Ken K. Ong, Ieuan A. Hughes, Carlo L. Acerini, David B. Dunger

## Abstract

**Aims:** The incidence of gestational diabetes has been reported to have risen over the first decade of this century. Some studies have also found it to vary with seasons of the year. We therefore investigated temporal and seasonal trends on gestational diabetes incidence in a single centre cohort study from Cambridge, U.K., and attempted to explain trends using associations with risk factors.

**Materials and Methods:** Using a cosinor model we tested whether there were both temporal and seasonal trends in gestational diabetes incidence in 1,074 women recruited to the Cambridge Baby Growth Study in 2001-2009 who underwent oral glucose tolerance tests around week 28 of pregnancy. We also undertook risk factor analyses.

**Results:** There was a temporal increase in gestational diabetes incidence over the course of recruitment to this study (p=2.1×10^−3^) but no seasonal effect (p=0.7). HOMA B (p=3.0×10^−3^; n=1,049) and the insulin disposition index (p=3.0×10^−3^; n=1,000) showed negative temporal trends. There was no negative association with HOMA S. Risk factor analyses showed a concomitant temporal slight increase in the index of multiple deprivation (p=4.6×10^−10^, n=1,068). This index was positively associated with HOMA B (p=6.1×10^−5^, n=955) but not directly with gestational diabetes (p=0.6, n=1,032), HOMA S (p=0.2, n=955) or the insulin disposition index (p=0.4, n=955).

**Conclusions:** In this population there were temporal but not seasonal increases in gestational diabetes incidence between the years 2001 and 2009, which appeared to be related more to reductions in insulin secretion than sensitivity. Possible mediators of this link include confounding factors related to deprivation.

## Introduction

Gestational diabetes (GDM) is traditionally defined as carbohydrate intolerance with its onset or first recognition in pregnancy,^1^ although more recent definitions explicitly exclude pre-existing type 2 diabetes.^2^ It is one of the most common adverse conditions of pregnancy. Its incidence has generally been reported to be rising in most populations, usually in line with the increasing prevalence of maternal obesity.^3^ Other major risk factors for GDM include having a previous history of it or having previously given birth to a macrosomic baby, a family history of GDM and/or type 2 diabetes, increased maternal age, increased gestational weight gain, genetics, multiparity and ethnic factors,^4^ not all of which can explain the tempo of the rising incidence. As GDM increases the risk of short- and long-term adverse complications for both the mother and her unborn child (including macrosomia, pre-eclampsia, childhood obesity and the metabolic syndrome in the mother^5^) and may contribute to the diabetes pandemic^6^, a thorough understanding of its pathogenesis is essential.

With the apparent worldwide rise in the prevalence of GDM^7^ paralleling the increasing prevalence of female obesity^8^ of note are the temporal and potentially mechanistic links that obesity has with global warming.^9^ Therefore another factor that could explain at least part of the increased incidence of GDM is exposure to raised or rising ambient temperatures in certain populations.^10-12^ Following this some studies have reported seasonal variations in the incidence of GDM^13-17^, although this has not been observed in all populations or climates.^18, 19^ In this study we investigated whether there were temporal and seasonal trends in GDM incidence in our single centre population from Cambridge, U.K. which recruited pregnant women between 2001 and 2009. We then investigated what may have mediated any trends. Although relating to a decade ago this seemed reasonable given that GDM was already becoming more prevalent by this time in several different populations.^20-25^

## Materials and Methods

### Cambridge Baby Growth Study

The prospective, longitudinal Cambridge Baby Growth Study (CBGS) was established as an observational cohort initially covering pregnancy, birth and infancy.^26^ 2,229 mothers, all over 16 years of age, were recruited when attending early pregnancy ultrasound clinics at the Rosie Maternity Hospital, Cambridge, U.K. between April 2001 and March 2009. 571 of these mothers withdrew prior to the birth of their infant. Most of the clinical characteristics of the study participants were collected either during nurse-led interviews or by questionnaire with the exception of offspring birth weight, gestational age and date of birth, which were compiled from hospital notes. In this cohort 95.3% of the offspring were white, 1.7% were Asian, 1.3% were black (African or Caribbean), and 1.7% were other ethnicities (mainly mixed race), reflective of the population served by the Rosie Maternity Hospital.

### Ethics

The Cambridge Baby Growth Study was approved by the Cambridge Local Research Ethics Committee, Addenbrooke’s Hospital, Cambridge, United Kingdom (00/325). All procedures followed were in accordance with the institutional guidelines. Written informed consent was obtained from all the study participants.

### Oral Glucose Tolerance Test and Gestational Diabetes Diagnostic Criteria

At a median (inter-quartile range) of 28.4 (28.1–28.7) weeks gestation 1,074 of the CBGS mothers underwent a 75g oral glucose tolerance test (OGTT) after fasting overnight.^27^ Venous blood was collected just prior to and 60 min. after the glucose load was administered for the measurement of plasma glucose, insulin and c-peptide concentrations. 120 min. plasma glucose concentrations were only measured from May 2007 onwards so were not used in this study to define GDM (only 7% of U.K. women with GDM receive a diagnosis based solely on the 120 min. measurement in any case^28^). The International Association of Diabetes in Pregnancy Study Groups (IADPSG) thresholds for 0 and 60 min. OGTT glucose concentrations (i.e. ≥ 5.1 and 10.0 mmol/L, respectively^29^) were used to define the presence of GDM.

### Assays

All biochemical kit-based assays were run according to the manufacturer’s instructions. Glucose concentrations were measured using a routine glucose oxidase-based method. OGTT plasma insulin concentrations were measured by ELISA (Dako UK Ltd., Ely, Cambs, U.K.). Intra-assay imprecision (CV) was 4.3% at 14 mU/L (82 pmol/L), 3.0% at 67 mU/L (402 pmol/L), and 5.7% at 151 mU/L (907 pmol/L). Equivalent inter-assay imprecision was 4.3, 5.1, and 5.4%, respectively. C-peptide concentrations were also measured by ELISA (DSL Labs., London, U.K.). Intra-assay imprecision was 2.8% at 1.3 ng/mL (0.43 nmol/L) and at 4.4 ng/mL (1.47 nmol/L), and 3.2% at 8.4 ng/mL (2.80 nmol/L). Equivalent inter-assay imprecision was 15.7%, 7.8% and 10.3%, respectively.

### Calculations

The maternal body mass index (BMI) was calculated as the pre-pregnancy weight divided by the height squared. Insulin sensitivity and pancreatic β-cell function were estimated using the homeostasis model assessment (HOMA S and B, respectively), calculated using the week 28 fasting circulating glucose and insulin (or c-peptide) concentrations and the online HOMA2 calculator (available at https://www.dtu.ox.ac.uk/homacalculator/).^30^ For this study HOMA values were calculated using both insulin and c-peptide concentrations separately, and where mentioned in this manuscript refer to insulin-derived values unless stated otherwise. Insulin secretion (for a given insulin sensitivity) was assessed in terms of the insulin disposition index, calculated as the change in insulin concentrations over the first hour of the OGTT divided by the change in glucose concentrations, all divided by the reciprocal of the fasting insulin concentration. An equivalent disposition index was also calculated using plasma c-peptide concentrations. The index of multiple deprivation was derived and imputed from the postcode of the participants’ home addresses as described.^31^

### Statistical Analysis

The present analysis was restricted to those 1,074 pregnancies where the women underwent OGTTs (thereby excluding women with pre-existing type 1 diabetes) with 0 and 60 min. plasma glucose concentrations available to us. Data were logarithmically transformed prior to analyses if their distributions were positively skewed. Temporal trends, adjusted for seasonal trends, were tested for using the cosinor regression model in R (version 3.5.2; The R Project for Statistical Computing, Vienna, Austria) deploying the package Cosinor (version 1.1, available at http://github.com/sachsmc/cosinor) which assumes a sinusoidal seasonal pattern over a longitudinal period, in this case OGTT season and year of analysis, respectively. The season was based on the month the OGTT was performed using Northern Meteorological seasons: winter (December to February), spring (March to May), summer (June to August), autumn (fall) (September to November). Further analysis was performed by standard logistic (for binary variables) or linear (for continuous variables whose model residuals using untransformed or transformed data were normally distributed) regression. Non-parametric regression (to fit linear regression models that included the index of multiple deprivation due to the lack of normal distribution of the residuals in standard linear regression models) was performed by means of the Theil–Sen procedure (deploying the R package mblm, version 0.12.1, available at https://cran.r-project.org/web/packages/mblm/index.html). Categorical analysis was performed using the χ^2^ test. Unless stated all the other statistical analyses were performed using Stata (version 13.1; Stata Corp., from Timberlake Consultants Ltd., Richmond, Surrey, U.K.). Statistical significance was assumed at p<0.05 throughout.

## Results

### Characteristics of the Study Population

With the exception of a slight increase in parity and a reduced proportion of smokers, those women who were included in the analysis were representative of the Cambridge Baby Growth Study as a whole (Table 1). Variables that were not detectably different included GDM prevalence, fasting glucose and insulin concentrations, pre-pregnancy BMI and maternal age.

**Table 1.**
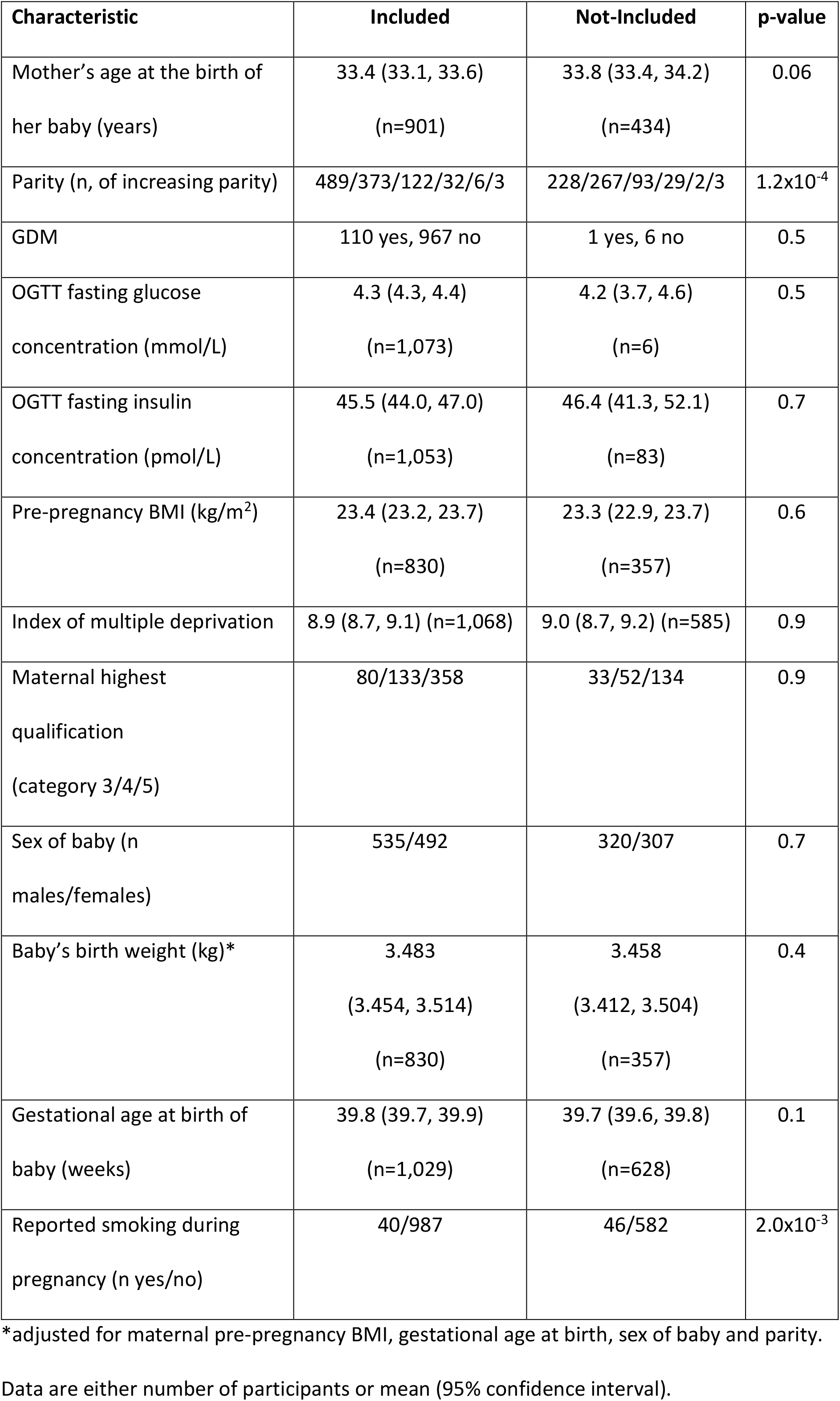
Characteristics of those Cambridge Baby Growth Study participants who were included in the current analysis and those that were not.

### Temporal Trends of GDM Incidence and OGTT Glucose Concentrations

The overall prevalence of GDM in this population was 10.1% (108/1,071). Cosinor analysis showed a significant temporal effect associated with the year of analysis (0.014 (0.005, 0.022) proportional increase per year, p=2.1×10^−3^), with a trend for the incidence of GDM increasing through the years 2001-9 (Fig. 1). This finding was confirmed by logistic regression (odds ratio (OR) 1.2 (1.1, 1.3) per year, p=2.8×10^−3^, n=1,049), even when the small numbers of participants collected in 2009 were excluded from the analysis (OR 1.2 (1.1, 1.3) per year, p=3.3×10^−3^, n=1,040). There was, however, no significant association of GDM with seasonality (amplitude: 9.8 (−14.7, 34.3), p=0.4; acrophase −0.6 (−3.0, 1.9), p=0.7) (Fig. 1). Cosinor analysis revealed no significant association of the OGTT fasting glucose concentration with the year (0.009 (−0.007, 0.025) mmol/L per year, p=0.3) or with seasonality (amplitude: 21.3 (−24.1, 66.6), p=0.4; acrophase −0.4 (−2.4, 1.6), p=0.7). Conversely there was a temporal trend with the OGTT 60 min. glucose concentrations (0.117 (0.067, 0.166) mmol/L per year, p<1×10^−4^) (Fig. 2), a finding which was confirmed by linear regression (β’=0.147, p=1.3×10^−6^, n=1,071), without a seasonal effect (amplitude: 53.7 (−84.1, 191.5), p=0.4; acrophase −0.9 (−3.4, 1.7), p=0.5).

**Fig. 1.**
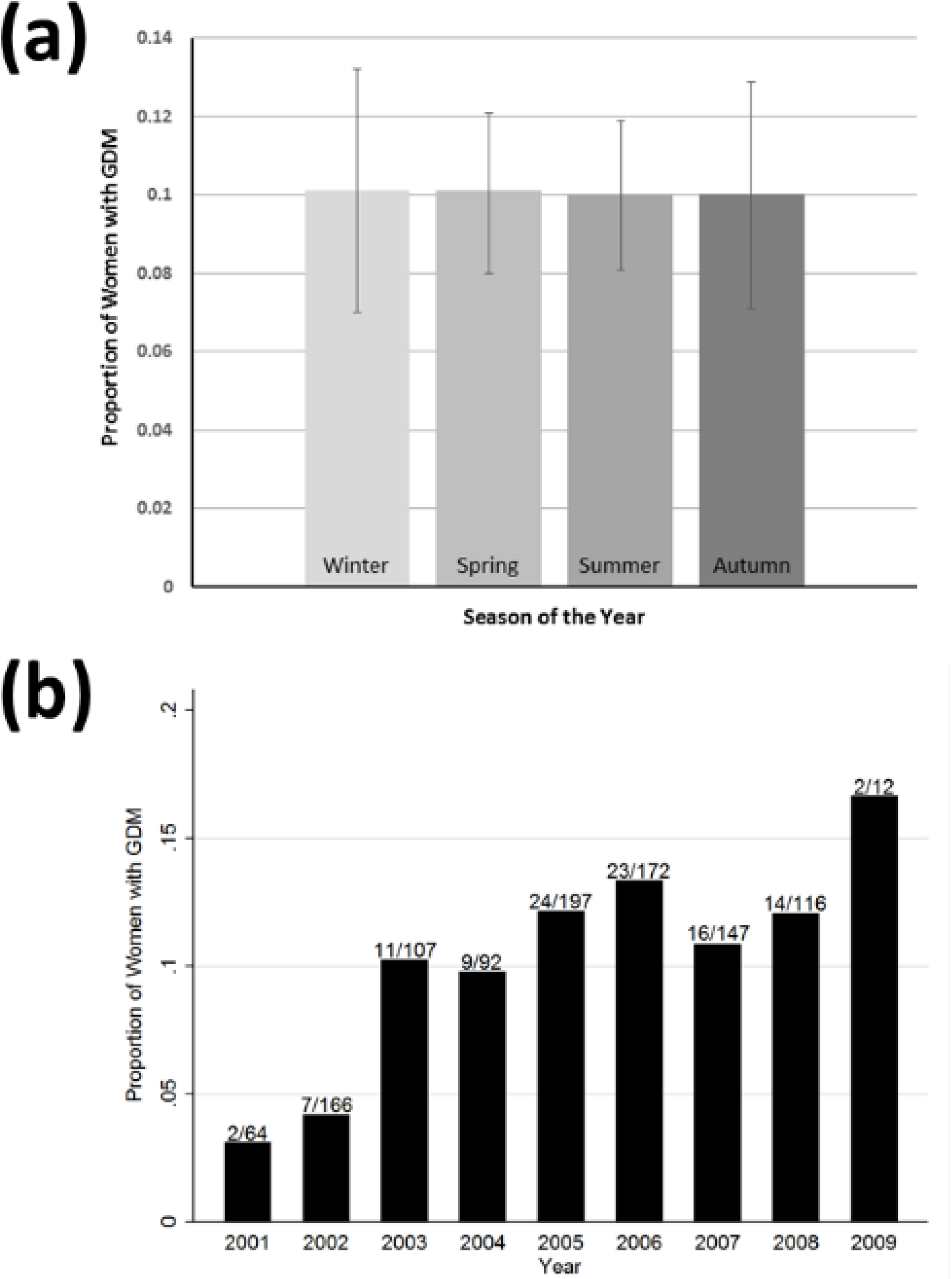
(a) The proportion of women with GDM by the season of the year, following an OGTT around week 28 of pregnancy, adjusted for year of analysis. Data are means (95% confidence intervals). Analysing this data categorically (taking no account of the recurring order of the seasons) showed no association between GDM and season of the year (χ^2^=2.2, p=0.5). (b) The proportion of women with GDM by the year of testing, following an OGTT around week 28 of pregnancy.

**Fig. 2.**
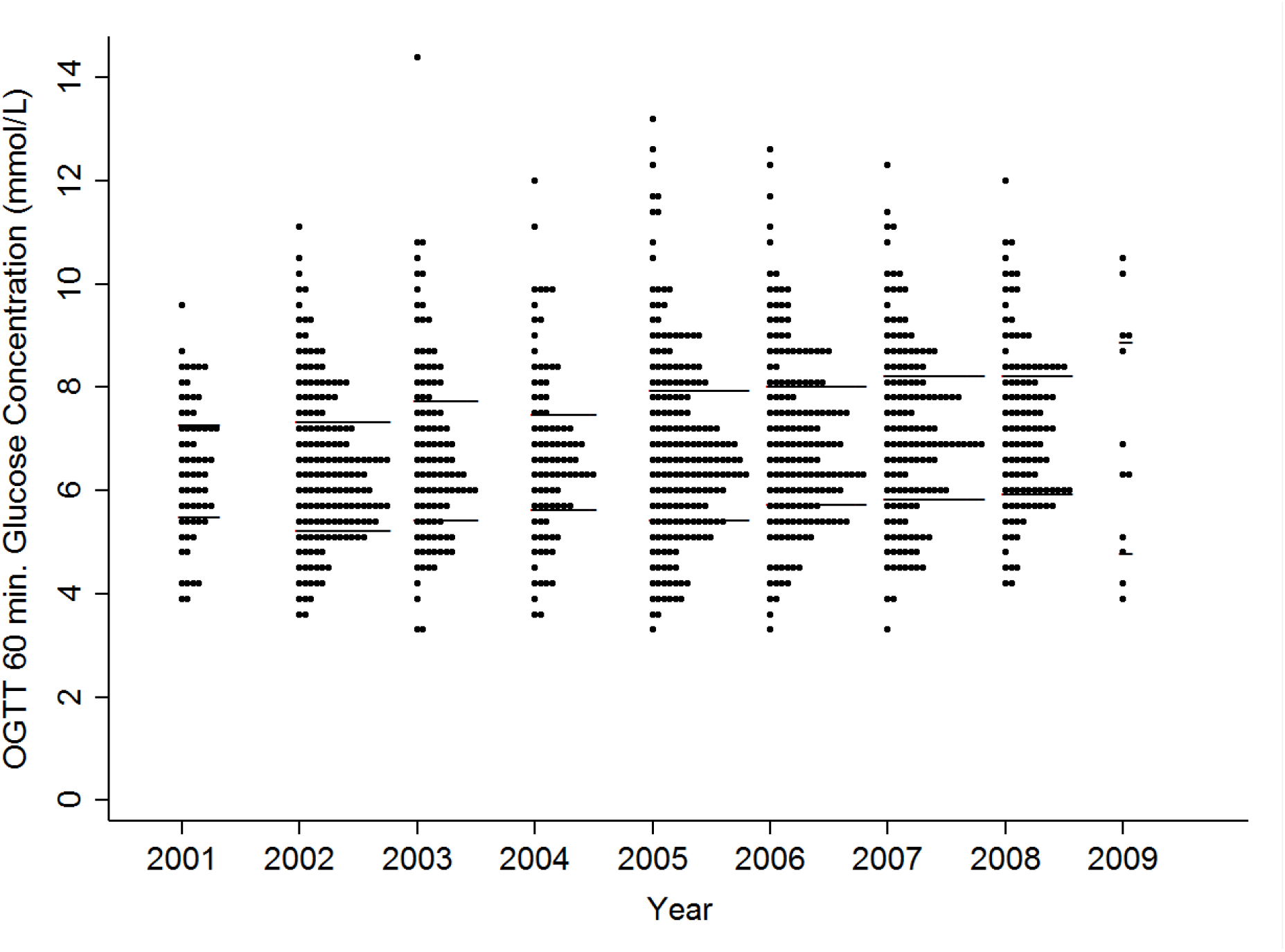
A dot plot of OGTT 60 min. glucose concentrations by the year of OGTT testing. The bars represent the limits of the interquartile range of the values represented by the dots.

### Temporal Trends of Indices of Insulin Sensitivity and Secretion

Insulin-derived HOMA S was positively associated with the year of testing (0.016 (0.001, 0.032) per year, p=0.04) but not with the season of testing (amplitude: 21.7 (−20.8, 64.1), p=0.3; acrophase 1.3 (−0.7, 3.4), p=0.2), a finding which was confirmed by linear regression (β’=0.080, p=9.1×10^−3^, n=1,049). This association was still evident if adjusting for GDM (p=3.4×10^−4^, n=1,049) or by only including women who were classified as not having GDM (p=1.2×10^−3^; n=947). However the association disappeared if it were tested using c-peptide-derived HOMA S, in a smaller number of women (β’=0.014, p=0.7, n=929).

Whilst still having a lack of association with seasonality (amplitude: 20.9 (−6.4, 48.2), p=0.1; acrophase 0.6 (−0.7, 1.9), p=0.4), in contrast to the findings with HOMA S, insulin-derived HOMA B was negatively associated with year of testing (−0.015 (−0.025, −0.005) per year, p=3.0×10^−3^). This association was confirmed by linear regression (β’=−0.104, p=7.0×10^−4^, n=1,049). Again it was still evident if adjusting for GDM (p=1.2×10^−3^, n=1,049) or if only including women without GDM (p=1.2×10^−3^; n=947). This association was still present if it were tested using c-peptide-derived HOMA B, in a smaller number of women (β’=−0.073, p=0.03, n=929).

Similar to the results for HOMA B, the insulin disposition index was negatively associated with year of testing (−0.036 (−0.060, −0.013) per year, p=3.0×10^−3^), a finding which was confirmed by linear regression (β’=−0.091, p=3.8×10^−3^, n=1,000). This association was still evident if adjusting for GDM (p=0.03; n=1,000) or if only including women without GDM (p=0.03; n=905). The insulin disposition index was not, however, associated with season of testing (amplitude: 26.0 (−38.5, 90.5), p=0.4; acrophase 1.1 (−1.5, 3.7), p=0.4). The c-peptide disposition index was also negatively associated with year of testing (β’=−0.074, p=0.03, n=882).

### Associations of the Year of Analysis with Potential Risk Factors

The year of analysis was not associated with either the maternal BMI (β’=0.006, p=0.9, n=827) or pregnancy weight gain (β’=−0.047, p=0.2, n=614). Neither was it associated with maternal age (β’=−0.016, p=0.6, n=898) or parity (β’=0.007, p=0.8, n=1,022) in this population. There was no association between the year of analysis and the proportion of male babies (χ^2^=4.9, p=0.8, n=1,024), birth weight of the babies (β’=−0.023, p=0.4, n=824; adjusted for gestational age at birth, mother’s BMI, parity and sex) or the odds of a pregnancy being multifetal (OR 1.0 (0.9, 1.2), p=0.9, n=1,071). One potential confounder that year of analysis was modestly albeit highly significantly associated with, the index of multiple deprivation (p=4.6×10^−10^, n=1,068) (Fig. 3), was itself not directly associated with GDM (OR 1.0 (1.0, 1.1) per unit increase in the index, p=0.6, n=1,032). There was still a significant relationship between the index of multiple deprivation and the year in the reduced number of 955 women for whom HOMA modelling was available (p=5.2×10^−8^). In these women the index of multiple deprivation was significantly positively associated with HOMA B (p=6.1×10^−5^; Fig. 3) but not with HOMA S (p=0.2) or the insulin disposition index (p=0.4).

**Fig. 3.**
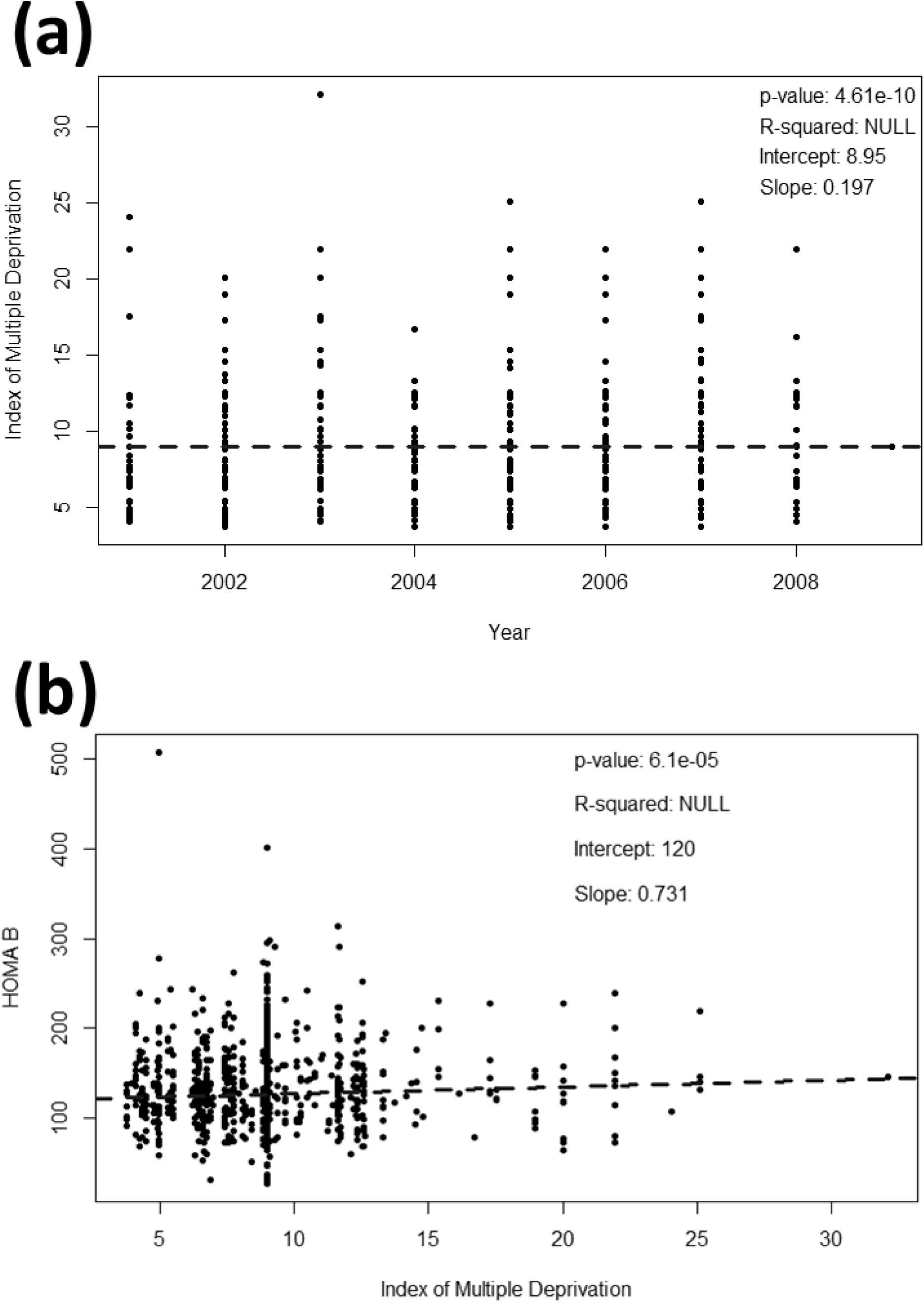
Dot plots of (a) the index of multiple deprivation by year of testing and (b) HOMA B values by the index of multiple deprivation. The lines of best fit (minimising the distance between the line and median values) were calculated using the Theil–Sen procedure.

## Discussion

In this analysis there was a strong trend for the incidence of GDM increasing as recruitment to the cohort progressed between 2001 and 2009. Trends over a similar period of time have also been observed in populations in Canada^21^, the U.S.A.^22, 24^, Israel^23^ and Germany^25^. The worldwide increased incidence at that time appeared to be independent of ethnicity^20^ despite this being a major factor associated with GDM risk.^4^ In our population the increased incidence of GDM was associated with reductions in HOMA B and the insulin disposition index (rather than HOMA S). Whilst a temporal trend in these factors could relate to changes in the performance of the insulin and glucose assays that were used over time, we used the same assays for these analytes throughout this time period and the performance characteristics of the insulin assay matched those of similar assays available at the time.^32^ Given that the performance of the glucose assay did not change over this time, the fact that the associations with HOMA B and the disposition index persisted if they were calculated using c-peptide rather than insulin concentrations suggests that these associations were physiological rather than assay-related. The temporal increase in GDM incidence in our population therefore appeared to relate to reductions in insulin secretion rather than sensitivity (indeed insulin-derived HOMA S actually increased over the period of recruitment, albeit this association was not evident when c-peptide concentrations were used to calculate HOMA S in a smaller number of women).

The temporal increase in GDM incidence was clearly environmentally-mediated as its tempo was too fast for a genetic change. In investigating its potential causes and the reduced insulin secretion we could not find parallel increases in BMI, pregnancy weight gain, maternal age or smoking. There was, however, a significant temporal trend of increasing deprivation in the study (in a population that was generally less-deprived than the national average^26^) although deprivation itself was not associated with GDM. There was a modest albeit highly significant positive association between the index of multiple deprivation and insulin-derived HOMA B. This suggests that, in the absence of a direct associations with HOMA S or GDM risk, as the deprivation index went up pancreatic β-cell function might have had to increase slightly to maintain glucose homeostasis. Due to the lack of direct association between the deprivation index and GDM despite the temporal trend, it is possible however that as yet unidentified confounder(s) related to deprivation, such as factors connected with diet^33^ and/or exercise^34^, could have contributed to the temporal increase in GDM incidence. This would be consistent with studies where associations between GDM and deprivation, lower socioeconomic status or lower social class were reported^25, 35, 36^ although such associations have not been found in every study.^37^ An alternative explanation for the temporal trend in deprivation in the present study is that it may just reflect unintentional secular patterning in study recruitment or uptake.

Although we observed a temporal trend of increasing GDM incidence as the decade progressed in this analysis we could not find a seasonal trend. This is despite seasonal trends in GDM previously have been observed in populations in Sweden^13^, Australia^14, 15^, Italy^16^ and Greece^17^. However of the populations tested before where no such trend was observed^18, 19^ one of these was also in the U.K.^19^ so our lack of seasonal trend may relate to climate or other environmental factors specific to the U.K. Alternatively, whilst at least one of the studies that found a seasonal trend used a very similar analysis technique to the one that we used^15^, other studies used analysis of variance or categorical/ordinal analyses which did not account for the recurring nature of the seasons or adjust for longer-term temporal trends^13, 14, 16, 17^ so differences from our results may relate to this.

The strengths of our prospective study include the fact that we used cosinor analysis to adjust linear temporal trends for separate potential seasonal effects, unlike some of the other studies in this area. In addition we had third trimester OGTT data from all the study participants and so we were able to investigate whether temporal trends in GDM incidence were related to changes in insulin sensitivity or secretion. We calculated HOMAs using both insulin and c-peptide concentrations so that detectable temporal trends were less likely to have resulted from drift in assay performance. In addition to its strengths the study does have a number of limitations however. Firstly this Cambridge cohort may not fully reflect the U.K. population as a whole particularly in relation to ethnic mix and smoking prevalence^25^, although this means that there was probably less confounding due to ethnic effects. Secondly we did not record family histories of GDM and type 2 diabetes, major risk factors for GDM.^4^ Thirdly although the study was fairly large given the level of detail that was collected, it was smaller than temporal studies of large surveys (e.g. ^38^) and therefore our trends were estimated with less precision than is possible in that kind of study. This could partially account for the large magnitude of the increased GDM incidence over the course of the recruitment period in our study. Finally the insulin (and c-peptide) disposition index was not calculated using 30 min. OGTT glucose and insulin concentrations like usual, but using 60 min. concentrations instead. Although comparing OGTT results from our study with those more commonly from intravenous GTTs, its use has been deemed acceptable at least for the insulinogenic element of the disposition index.^39^

In conclusion we observed a temporal but not seasonal trend for an increasing incidence of GDM in Cambridge Baby Growth Study pregnancies from the years 2001-2009. This was associated with reductions in indices of insulin secretion rather than insulin sensitivity. Although we do not know what caused these changes it does not appear to relate to changes in maternal obesity or age. Factors relating to deprivation offer potential explanations.

## Acknowledgments

The authors would like to thank all the families that took part in the Cambridge Baby Growth Study, and acknowledge the crucial role played by the research nurses especially Suzanne Smith, Ann-Marie Wardell and Karen Forbes, staff at the Addenbrooke’s Wellcome Trust Clinical Research Facility, and midwives at the Rosie Maternity Hospital in collecting data for this study. We would also like to thank Angie Watts, Karen Whitehead and Dianne Wingate for excellent laboratory assistance. This work was supported by the Medical Research Council (grant numbers G1001995, 7500001180); European Union Framework 5 (grant number QLK4-1999-01422); the Mothercare Charitable Foundation (grant number RG54608); Newlife – The Charity for Disabled Children (grant number 07/20); the World Cancer Research Fund International (grant number 2004/03); and the National Institute for Health Research Cambridge Biomedical Research Centre. KO is supported by the Medical Research Council (Unit Programme number: MC_UU_12015/2). None of the funding bodies influenced the design of the study or the collection, analysis or the interpretation of the data used in this study. Neither did they influence the writing of this manuscript.

## Author’s Contribution Statement

CP conceived and designed the study, analysed some of the data and wrote the initial draft of the manuscript. BF analysed some of the data and commented on the initial draft of the manuscript. KO, IH, CA and DD designed, established and oversee the Cambridge Baby Growth Study. All authors commented on early drafts of the manuscript and approved the submission of the final draft.

